# Impact of ploidy and pathogen life cycle on resistance durability

**DOI:** 10.1101/2021.05.28.446112

**Authors:** Méline Saubin, Stéphane De Mita, Xujia Zhu, Bruno Sudret, Fabien Halkett

## Abstract

The breeding of resistant hosts based on the gene-for-gene interaction is crucial to address epidemics of plant pathogens in agroecosystems. Resistant host deployment strategies are developed and studied worldwide to decrease the probability of resistance breakdown and increase the resistance durability in various pathosystems. A major component of deployment strategies is the proportion of resistant hosts in the landscape. However, the impact of this proportion on resistance durability remains unclear for diploid pathogens with complex life cycles. In this study, we modelled pathogen population dynamics and genetic evolution at the virulence locus to assess the impact of the ploidy (haploid or diploid) and the pathogen’s life cycle (with or without host alternation) on resistance durability. Ploidy has a strong impact on evolutionary trajectories, with much greater stochasticity and delayed times of resistance breakdown for diploids. This result emphasises the importance of genetic drift in this system: as the virulent allele is recessive, positive selection on resistant hosts only applies to homozygous (virulent) individuals, which may lead to population collapse at low frequencies of the virulent allele. We also observed differences in the effect of host deployment depending on the pathogen’s life cycle. With host alternation, the probability that the pathogen population collapses strongly increases with the proportion of resistant hosts in the landscape. Therefore, resistance breakdown events occurring at high proportions of resistant hosts frequently amount to evolutionary rescue. Last, life cycles correspond to two selection regimes: without host alternation (soft selection) the resistance breakdown is mainly driven by the migration rate. Conversely, host alternation (hard selection) resembles an all-or-nothing game, with stochastic trajectories caused by the recurrent allele redistributions on the alternate host.

## Introduction

Plant pathogens can quickly evolve (Perkins et al., 2013), and the loss of host genetic diversity in agroecosystems compared to natural ecosystems can enhance the spread of epidemics (Burdon and Thrall, 2008; Garrett et al., 2009; Haas et al., 2011; Mundt, 2002; Ostfeld and Keesing, 2012; Zhan et al., 2015). In this context, many plant protection strategies are developed and studied worldwide (Bousset and Chèvre, 2013), particularly spatio-temporal host resistance deployment strategies (Bousset et al., 2018; Burdon et al., 2014; Djian-Caporalino et al., 2014; Fabre et al., 2015; Gilligan and Bosch, 2008; Mundt, 2002; Rimbaud et al., 2018; Sapoukhina et al., 2009). However these modelling studies seldom account for pathogen differences in life cycle and ploidy levels.

While quantitative resistance has gained interest (Pilet-Nayel et al., 2017), the breeding of disease resistant plants is still often based on the gene-for-gene interaction (Person et al., 1962; Zhan et al., 2015). In the simplest case of specific response, the result of the infection is determined by the interaction between a locus in the plant (the resistance gene) and in the pathogen (the avirulence gene) (Flor, 1971). This interaction leads to an all-or-nothing response and therefore such resistances are called qualitative. Qualitative resistances often rely on the recognition of a specific pathogen molecule (an effector protein for instance) by a plant immune receptor (Lo Presti et al., 2015). If the pathogen is recognised by the plant, the infection is stopped and the plant is called *resistant*. But the pathogen species evolves in multiple ways to escape host recognition (Rouxel and Balesdent, 2017). When a pathogen can infect a resistant host it is called *virulent*, as opposed to *avirulent* individuals. For avirulent individuals, if the product of the avirulence gene is not recognised by the plant, the infection occurs and the plant is called *susceptible*. Hence, virulent individuals can infect both susceptible and resistant hosts, while avirulent individuals can only infect susceptible hosts. In its simplest cases, the avirulence gene exists in two versions: the avirulent *Avr* allele and the virulent *avr* allele. The plant resistance can thus be overcome by a mutation of the *Avr* allele which modifies the pathogen recognition by the plant. The *Avr* allele is then replaced by a virulent *avr* allele which leads to a virulent pathogen (Stukenbrock and McDonald, 2009).

In natural systems, the constant turnover of resistance and avirulence genes results from a strong coevolutionary interaction between both species (Zhan et al., 2014), represented by the concept of arms-race (Brown and Tellier, 2011). On both sides, the most adapted allele can spread in the population, sometimes replacing alleles conferring lower fitness to individuals (Brown and Tellier, 2011; Persoons et al., 2017). These genes are under strong selective pressure and at each selective event a selective sweep can occur and drastically reduce the genetic diversity of both species (Oleksyk et al., 2010; Terauchi and Yoshida, 2010). In natural populations, rare host genotypes can be maintained by negative frequency-dependent selection, resulting in the preservation of host polymorphism (Lewontin, 1958). In agroecosystems, however, pure crops of resistant hosts hinder this maintenance of polymorphism (Zhan et al., 2015). Therefore, the issue of such resistance deployments is often a resistance breakdown, *i*.*e*. the failing of the host to remain resistant to the pathogen, which can result in severe epidemics (Brown and Tellier, 2011; Burdon et al., 2016; Johnson, 1984; Pink and Puddephat, 1999). Such a resistance breakdown can occur more or less quickly, depending on the pathosystem considered and the environmental conditions (Van den Bosch and Gilligan, 2003). This observation raises the question of resistance durability, which can be defined as the time until the virulent population reaches a given threshold in the pathogen population. Definitions of resistance durability can have diverse acceptations depending on the threshold considered (Brown, 2015; Carolan et al., 2017; Lof et al., 2017; Pacilly et al., 2018; Pietravalle et al., 2006; Rimbaud et al., 2021; Van den Bosch and Gilligan, 2003). Considering several thresholds can help in capturing different steps of the pathogen dynamics.

Resistance durability becomes a major economical issue when epidemics impact crop yields. Therefore, it has often been studied through the modelling of epidemics spread in agricultural landscapes (Rimbaud et al., 2021). Such models can couple epidemiological and evolutionary processes, and often aim to study the influence of different biological parameters on the emergence of pathogens, their specialisation to the host plant, the evolutionary dynamics of virulence, or on the resistance durability (Van den Bosch and Gilligan, 2003). *Virulence* is defined here as the ability for a pathogen individual to infect a resistant host, in accordance to the phytopathology literature (Flor, 1971; McDonald and Linde, 2002). These parameters can be specific to the host plant (proportion of resistant host in the landscape, their spatial and temporal distribution) or to the pathogen (life cycle, mutation rate, dispersal) (Fabre et al., 2015, 2012; Papaïx et al., 2015, 2018; Soularue et al., 2017). These models often represent haploid pathogens with a virulent and an avirulent genotype, evolving purely asexually on a landscape composed of compartments, gathering resistant or susceptible hosts (Lof et al., 2017; Lof and Werf, 2017; Pacilly et al., 2018; Pietravalle et al., 2006). Regarding the pathogen, high risks of resistance breakdown are observed for pathogen populations with high gene flow and mutation rates, large effective population sizes, and partially asexual reproductions (McDonald and Linde, 2002). Regarding the host, the increase in the proportion of resistant hosts should increase the selection pressure, hence weakening the resistance durability (Pietravalle et al., 2006; Van den Bosch and Gilligan, 2003). However, a large proportion of resistant hosts also reduces the initial size of the pathogen population and thus the risk of resistance breakdown (Pacilly et al., 2018), partly because a small population size reduces the likelihood that a virulent individual will emerge through mutation.

However, the impact of host resistance deployment on resistance durability remains unclear when the pathogen is diploid (like rust fungi, oomycetes, or nematodes). When the product of the avirulence gene is a specific molecule like an effector protein, the pathogen is virulent only if this product is not detected by the product of the corresponding resistance gene in the host (Stukenbrock and McDonald, 2009). Therefore, for a diploid individual the pathogen is virulent only if the products of both alleles avoid detection by the host. In other words, in the classical gene-for-gene interaction the virulent allele is recessive (Thrall et al., 2016). Consequently, a heterozygous *Avr*/*avr* individual is phenotypically avirulent, and the selective advantage of the virulence is effective among homozygous *avr*/*avr* individuals only. At low frequency, *avr* alleles are then carried by heterozygous individuals and mostly subjected to drift.

Diploid pathogens exhibit a large variability of life cycles (Agrios, 2005). We can especially distinguish autoecious pathogens, which complete their life cycle on a unique host species, from heteroecious pathogens which need two different and successive host species to complete their life cycle (Lorrain et al., 2019; Moran, 1992). This presence or absence of an alternate host species could also affect the influence of host deployment strategy on resistance durability. Moreover, most studies focus on purely asexual pathogens, but the highest risks of resistance breakdown were observed for mixed reproduction systems (McDonald and Linde, 2002), with the best invaders combining high rates of asexual reproduction and rare events of sex (Bazin et al., 2014). Yet, the allelic redistribution resulting from a sexual reproduction event could have an even stronger impact on resistance durability when the pathogen is diploid.

To study resistance durability and evolutionary forces shaping the system, the understanding of the evolution of gene and virulence allele frequencies is needed. Coupling epidemiology and population genetics provides insights on both short and long time scales. It allows in particular detailed analyses of transition periods (Bolker et al., 2010; Day and Gandon, 2007; Day and Proulx, 2004), through variables like the pathogen population size, affecting both the disease incidence in epidemiology and the impact of genetic drift in population genetics (McDonald, 2004). This approach is also crucial for highlighting transient effects like evolutionary rescue, *i*.*e*. the genetic adaptation of a population to a new environment, thus preventing its extinction (Alexander et al., 2014; Martin et al., 2013). Evolutionary rescue as a process leading to resistance breakdown has not received consideration so far.

The virulence of pathogens can be associated with a fitness cost on susceptible hosts (Bousset et al., 2018; Laine and Barrès, 2013; Leach et al., 2001; Montarry et al., 2010; Nilusmas et al., 2020; Thrall and Burdon, 2003), sometimes referred to as the cost of pathogenicity (Sacristán and García-Arenal, 2008). This fitness cost has been shown to have a strong impact on resistance durability (Fabre et al., 2012). However, depending on the avirulence gene considered, such a fitness cost is not systematic (see Leach et al., 2001 for a review). Therefore, in the absence of data, it could be more conservative of the risk of breakdown not to consider fitness cost while modelling resistance durability.

In this paper, we aim to broaden our understanding about the impact of the ploidy and the life cycle of pathogens on resistance durability. We used a non-spatialised model coupling population dynamics and population genetics to simulate the evolution of pathogens on susceptible and resistant hosts. We investigated the effects of resistant host deployment and pathogen demography on resistance durability, for a population of diploid and partially clonal pathogens, and compared the results to those obtained for haploid pathogens. Two different life cycles were implemented specifically: with or without host alternation for the sexual reproduction of the pathogen population. We assessed the resistance durability in two steps. First we examined the dynamics of fixation of the virulent allele in the population, and considered in parallel the cases when the pathogen population could go extinct, to highlight evolutionary rescue events. Then we focused on the invasion and resistance breakdown events, and disentangle the relationship between the durability of resistance and the dynamics of virulent populations after the invasion of the resistant plants.

## Model description

### Model overview

The model is individual-based, forward-time and non-spatialised, and couples population dynamics and population genetics to study the evolution of a population of pathogens for a succession of generations. It allows us to follow the evolutionary trajectory of different genotypes at the avirulence locus of pathogens through time. We consider a life cycle usually found in temperate pathogen species, which alternate rounds of clonal reproduction with an annual event of sexual reproduction (Agrios, 2005). This model is designed in four variants to represent haploid or diploid pathogens with two distinct life cycles: with or without host alternation for the sexual reproduction (Figure 1). Without host alternation, the model represents the evolution in time of a population of pathogens on two static host compartments: a resistant compartment (R) and a susceptible compartment (S). Fixed carrying capacities of pathogens, *K*_*R*_ and *K*_*S*_, are respectively assigned to R and S compartments and represent the maximum amount of pathogens that each compartment can contain. With host alternation, the alternate host compartment (A) is added, where the sexual reproduction takes place. This static compartment is assumed to be sufficiently large and thus with unbounded population size. Note that the life cycle with host alternation for haploid pathogens was added for the sake of comparison but has no real biological meaning, because no haploid pathogen display this life cycle.

**Figure 1.**
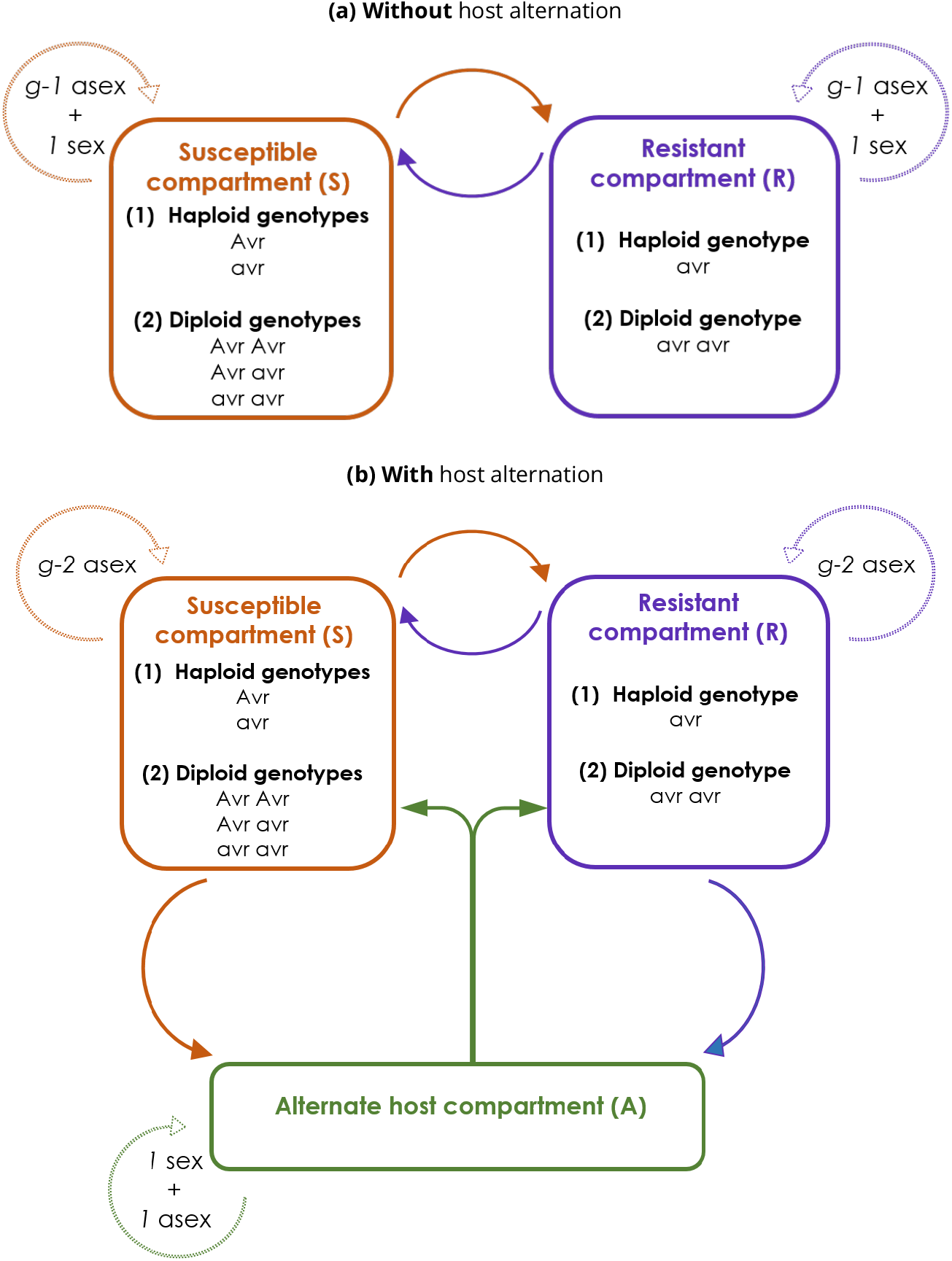
Model representation for two different life cycles: (a) without or (b) with host alternation. *g* corresponds to the total number of generations (asexual plus sexual) in a year. Dashed arrows represent reproduction events, and solid arrows represent migration events occurring at each generation. *asex* stands for asexual reproduction events, and *sex* for sexual reproduction events. *avr* denotes the virulent recessive allele, and *Avr* the avirulent dominant allele.

### Demographic evolution of the pathogen population

#### Reproduction events

Each discrete generation corresponds to a reproduction event, either sexual or asexual. Each year is composed of *g* non-overlapping generations, with one annual sexual reproduction event followed by a succession of *g* − 1 asexual reproduction events. In our simulations, we considered a year composed of *g* = 11 generations. At each reproduction event, parents give way to offspring and the new population is composed exclusively of new individuals. The within-compartment dynamics of the pathogen population are provided by the following equations:

Before each sexual reproduction event, a proportion *Reduct* of pathogen individuals is picked randomly to form the parental population. We fixed *Reduct* = 0.2 to cope to pathogen life cycles displaying drastic reduction in population size during sexual reproduction which usually takes place in winter.

For the sexual reproduction event itself, the population size is considered constant before and after the reproduction event:

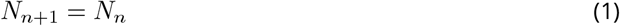

With *N*_*n*+1_ the population size at generation *n* + 1 and *N*_*n*_ the population size at generation *n*. Sexual offspring genotypes are obtained through random mating within the parental population.

For the asexual reproduction following the sexual reproduction in the A compartment in the life cycle with host alternation, the population growth is exponential, with the following relation:

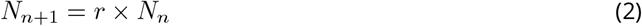

With *r* the growth rate of the pathogen population.

For each asexual reproduction in the R or S compartments, the population growth is logistic, with the following relation:

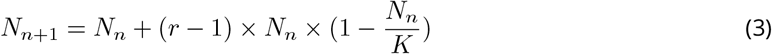

With *K* the carrying capacity of the compartment (*K*_*R*_ or *K*_*S*_ for R or S compartment respectively). For all clonal reproduction events, offspring genotypes are drawn randomly from the parental population, with replacement, considering the same reproduction rate for all pathogen genotypes.

#### Migration events

A regular two-way migration event takes place each generation before the reproduction event, between individuals evolving in the R and S compartments. The number of migrant individuals is determined by a migration rate (*mig*) multiplied by the number of individuals on the compartment of origin. Migrant individuals succeed to invade the compartment of arrival, even if the number of individuals on this compartment reached the maximum carrying capacity. The population size on each compartment is restricted to the carrying capacity during reproduction events only, and not during migration events. Thereby, this choice enables the immigration of new pathogens regardless of the size of the population, as it is observed in natural populations for plant pathogens.

For the life cycle with host alternation, the annual sexual reproduction event coincides with the obligate migration of the entire pathogen population to and from the alternate host. The first migration event takes place once every year after *g* − 2 asexual reproduction events in the R and S compartments (Figure 1). For this migration event, an established proportion of individuals *Reduct* is picked randomly from R and S compartments to migrate to the A compartment. All remaining individuals die in the R and S compartments, because the sexual reproduction is mandatory to complete the life cycle. After the two reproduction events (sexual and asexual) in the A compartment, the second migration event redistributes randomly all individuals from the compartment A to R and S compartments, in proportion to the relative size of R and S compartments (Figure 1).

### Genetic evolution of the pathogen population

To better highlight the effect of drift among other evolutionary forces, we did not consider mutation, that is, there is no change by chance of allelic state. This amounts to study evolution of the pathogen population from standing genetic variation (Barrett et al., 2008). The avirulence gene exists at two possible states: the *Avr* allele and the *avr* allele. For haploid pathogens, the *Avr* allele leads to avirulent individuals surviving only in the S compartment (and in the A compartment in the case of host alternation), while the *avr* allele leads to virulent ones capable to survive on all compartments without any fitness cost (Brown, 2015; Leach et al., 2001). For diploid pathogens, *Avr* is dominant and *avr* is recessive. Thus, individuals with genotypes *Avr*/*Avr, Avr*/*avr* and *avr*/*avr* survive with equal fitness in the S and A compartments, while only individuals with the virulent genotype *avr*/*avr* survive in the R compartment. Every avirulent individual (*Avr* for haploids, and *Avr*/*Avr* or *Avr*/*avr* for diploids) migrating to the R compartment dies before any subsequent migration or reproduction event.

Besides the demographic evolution of pathogen populations, the model describes the evolution of allelic and genotypic frequencies through generations in each compartment. Reproduction events can change allelic and genotypic frequencies. In particular, the annual sexual reproduction is the only event responsible for allele reshuffling in diploid individuals. For haploid pathogens, as only one locus is studied, the sexual reproduction event amounts to asexual reproduction, with differences in the size of the offspring population only.

Resistance durability is evaluated at four steps representing different proportions of virulent individuals in the population: (1) the time of apparition of the first virulent individual on the R compartment; (2) the time of invasion of the R compartment (1‰ of the R compartment occupied); (3) the time of resistance breakdown (1% of the R compartment occupied); and (4) the time of fixation of the virulence (all individuals are virulent, i.e only *avr* alleles remain). The thresholds of 1‰ and 1% were arbitrarily fixed to correspond to (i) the establishment of a pathogen population on the R compartment for the invasion and (ii) the detection of the virulent population on the R compartment for the resistance breakdown, respectively.

### Implementation of model analyses

The model was implemented in Python (version 3.7, Rossum, 1995), with the package “simuPOP” (Peng and Kimmel, 2005). The starting point of each replicate simulated was a population of 2000 individuals in the susceptible compartment. A proportion *f*_*avr*_ of virulent alleles was introduced initially in the pathogen population at Hardy-Weinberg equilibrium, as standing genetic variation. For diploid individuals, homozygous *avr*/*avr* individuals could therefore be initially present, depending on *f*_*avr*_. All simulations were run with a fixed total carrying capacity for the host population size, *K* = *K*_*R*_ + *K*_*S*_ = 100 000, but a variable proportion of the size of the R compartment 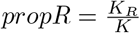

Preliminary analyses were carried out to study demographic and genetic outcomes with varying parameters. These analyses enabled six variables of interest to be identified: the initial frequency of *avr* allele (*f*_*avr*_), the migration rate (*mig*), the growth rate (*r*), the proportion of resistant hosts in the landscape (*propR*), the ploidy (*Ploidy*) and the life cycle (*Cycle*). Statistical analyses were performed on simulations with quantitative input parameters picked randomly from known distributions, resulting into a random simulation design (Table 1). The same simulation design was run four times, once for each combination of categorical input parameters (*Ploidy* and *Cycle*). To investigate further the impact of the input parameters on the simulation outcome in specific cases and to present the model results in a more didactic form, a regular simulation design was developed to complement the random design (Table 1).

**Table 1.** Input parameters and their range of variations for the three simulation designs. For both the random and regular simulation designs, 100 replicates were run for each combination to analyse the equilibrium reached by the population of pathogens after 1100 generations. For the restricted simulation design, 1000 replicates were run for each combination and the allele frequencies and population sizes were monitored through all 1100 generations simulated.

| Variable | Description | Random design |  | Regular design | Restricted design |
| --- | --- | --- | --- | --- | --- |
|  |  | Distribution | Interval |  |  |
| $f_{avr}$ | Initial frequency of the *avr* allele in the pathogen's population | Log-uniform | [log(0.0005), log(0.3)] | **6 levels:** 0.0005; 0.002; 0.01; 0.05; 0.15; 0.3 | 0.02 |
| *mig* | Migration rate between R and S compartments | Log-uniform | [log(0.001), log(0.2)] | **3 levels:** 0.05; 0.1; 0.2 | **2 levels:** 0.001; 0.05 |
| *r* | Growth rate of the pathogen | Uniform | [1.1, 2] | **10 levels :** 1.1; 1.2; 1.3; 1.4; 1.5; 1.6; 1.7; 1.8; 1.9; 2.0 | 1.5 |
| *propR* | Proportion of resistant hosts in the landscape | Uniform | ]0, 1[ | **9 levels:** 0.05; 0.1; 0.2; 0.35; 0.5; 0.65; 0.8; 0.9; 0.95 | **3 levels:** 0.1; 0.5; 0.9 |
| *Ploidy* | Ploidy of the pathogen | - | Haploid or Diploid | **2 levels:** Haploid; Diploid | **2 levels:** Haploid; Diploid |
| *Cycle* | Life cycle of the pathogen | - | Without or with host alternation | **2 levels:** Without host alternation; With host alternation | **2 levels:** Without host alternation; With host alternation |

This regular simulation design allowed us to present the results in a more conventional form. For both the random and the regular simulation design, simulations were run for each combination of parameters for 100 years (1100 generations) with 100 replicates. During this period, nearly all replicates reached equilibrium (fixation of one allele or extinction of the population).

A principal component analysis was performed on the data obtained with the random simulation design using R (R, 2018), on the following output variables: the frequency of extinction of population (*freq*_*ext*), the frequency of fixation of the *Avr* allele in the population (*freq*_*fix*_*Avr*), the frequency of fixation of the *avr* allele (*freq*_*fix*_*avr*) and the generation of fixation of *avr* (*gen*_*fix*_*avr*). To study the influence of the six input parameters (*Ploidy, Cycle, f*_*avr*_, *mig, r*, and *propR*) on the three main output variables selected (*freq*_*ext, freq*_*fix*_*avr*, and *gen*_*fix*_*avr*), generalized linear models (GLM) were performed on R. GLM on *freq*_*ext* and *freq*_*fix*_*avr* were performed with a Logistic link function, and the GLM on *gen*_*fix*_*avr* was performed with a Gamma link function.

To analyse further the temporal dynamics of *avr* allele frequency and population size, simulations were run recording population size and allelic states over time. Because these simulations were time- and memory-consuming, they were run on a restricted simulation design with only 24 combinations of parameters (Table 1). The generation of fixation of the *avr* allele was thus decomposed into two distinct output variables: the year of invasion of the R compartment and the time elapsed between the invasion and the *avr* allele fixation in the population. The influence of three parameters (*propR, Ploidy* and *Cycle*) on these two output variables was studied with 1000 replicates for each combination through 1100 generations. For these two output variables, GLM were performed with a Gamma link function.

For each general linear model developed, a dominance analysis was performed with the R package “dominanceanalysis” (Bustos Navarrete and Coutinho Soares, 2020) to compare the relative importance of the input parameters on the five output variables described. Estimated general dominance were calculated using bootstrap average values with the corresponding standard errors for each predictor with 100 bootstrap resamples, with McFadden’s indices (McFadden, 1974).

Calculations of a growth rate threshold *r*_0_ were carried out on Python for several parameter combinations. This value determines the growth rate below which the population goes extinct before the end of the simulation if there are no virulent individuals, therefore if the R compartment remains empty.

## Results

### Model behaviour

Since the model leads to stochastic outputs, we first analysed model behaviour in order to identify sound output variables. We visualised both population size and allele frequency dynamics through generations. The trajectory of each simulation either lead to the extinction of the entire pathogen population or to the fixation of one allele, provided that simulations last long enough. An example of such model behaviour is illustrated in Annex B (Figure S2) with four replicates, assuming diploid pathogens with host alternation. In this example, population sizes display cyclical dynamics due to the annual migration event on the A compartment. Three out of five replicates lead to population extinctions, while in the two other replicates, some pathogen individuals succeed to invade the R compartment after the initial phase of population dynamics collapse, leading to the fixation of the *avr* allele. These two dynamics with the survival of the population following a genetic adaptation to harsh environment illustrates evolutionary rescue. Interestingly, all replicates succeed to invade the R compartment at some time, but - because of host alternation - the annual redistribution of individuals breaks the invasion dynamics of the R compartment. Therefore, invasion does not necessarily lead to *avr* fixation.

In the following, we will summarise simulation results with four output variables, computed over replicates: the frequency of extinction, *freq*_*ext*; the frequency of fixation of *Avr* or *avr* allele, *freq*_*fix*_*Avr* or *freq*_*fix*_*avr*, respectively; and the generation of fixation of *avr* allele, *gen*_*fix*_*avr*. The later output variable provides insights on the durability of resistance.

### Sensitivity analyses

To identify the most significant parameters on the different output variables, we conducted a PCA analysis, general linear models and dominance analyses.

The PCA analysis was performed on the four output variables, with the first and second axes accounting for 49.5% and 33.3% of the total variability respectively (Figure 2). The most influential parameters of interest on the output variables were the growth rate *r*, negatively correlated with the frequency of extinction of population *freq*_*ext*. The initial frequency of *avr* alleles in the population *f*_*avr*_ was positively correlated with the frequency of fixation of the *avr* allele *freq*_*fix*_*avr*. The migration rate *mig* and the proportion of resistant hosts in the landscape *propR* were negatively correlated with both the frequency of fixation of the *Avr* allele *freq*_*fix*_*Avr*, and the generation of fixation of the *avr* allele *gen*_*fix*_*avr*. The qualitative input parameters (*Ploidy* and *Cycle*) were studied by representing each of the combinations of parameters of the random simulation design colored according to its ploidy and life cycle (Figure 2.b). This PCA highlights a higher frequency of extinctions of population for diploids with host alternation. Moreover, simulations without host alternation lead to higher frequencies of fixation of *Avr*, and longer times to *avr* fixation. The ellipses also illustrate that haploid individuals with host alternation lead to less variable outcomes and to higher frequencies of fixation of *avr*.

**Figure 2.**
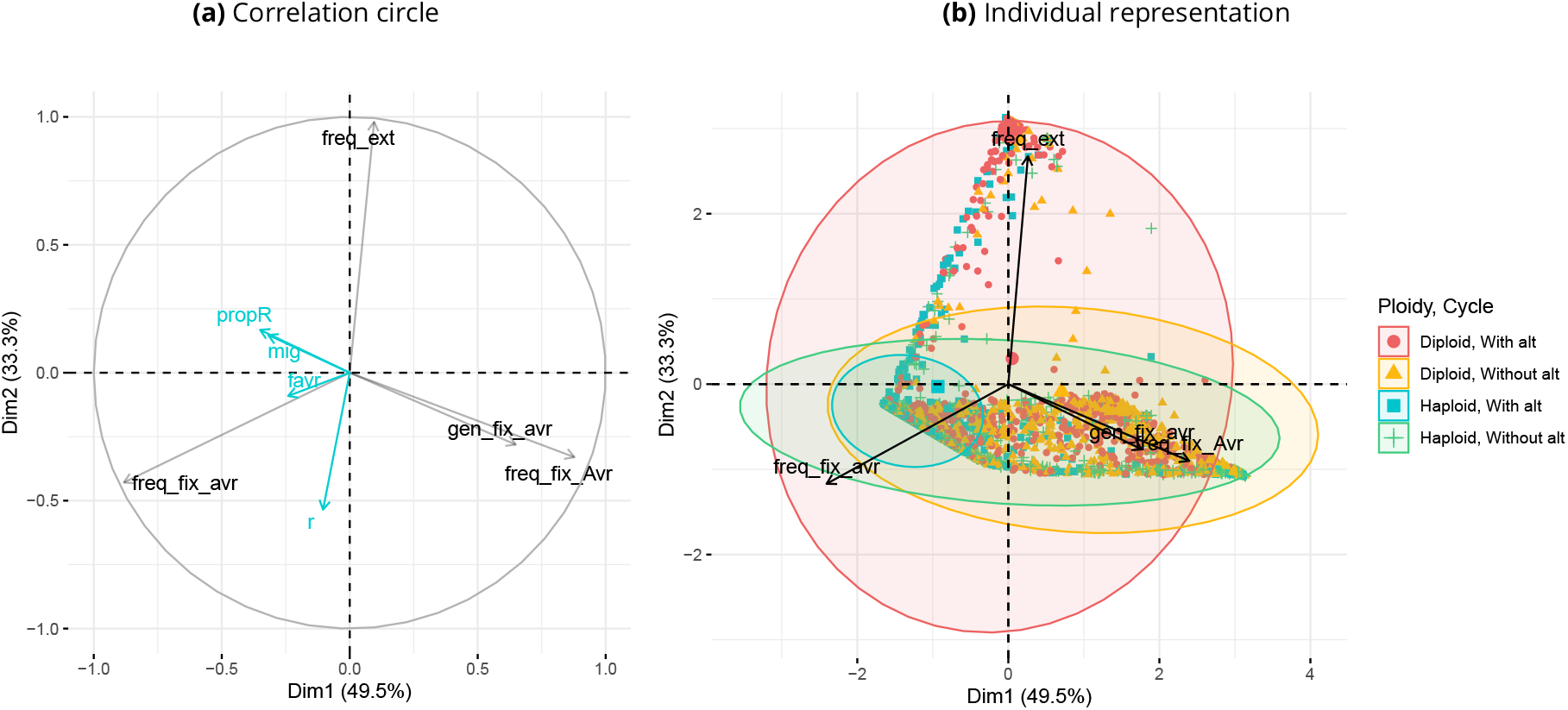
Principal component analysis on four output variables: *freq*_*ext, freq*_*fix*_*Avr, freq*_*fix*_*avr*, and *gen*_*fix*_*avr*. Two results are displayed: (a) the correlation circle on four output variables. The quantitative parameters *f*_*avr*_, *mig, r*, and *propR* are represented informatively in blue on the correlation circle and do not contribute to the variability; (b) the individual representation of simulations which represents the influence of the pathogen ploidy and life cycle. Each point represents a different combination of parameters with 100 replicates. Ellipses correspond to the 95% multivariate distribution.

Dominance analyses highlight the high impact of *r* on *freq*_*ext* (Figure 3.a), which is confirmed by the analysis of Sobol indices (Annex A Figure S1). For *freq*_*fix*_*avr* and *gen*_*fix*_*avr*, the influence of model parameters is more balanced with a lower contribution of the input parameters on *gen*_*fix*_*avr* (Figure 3.b, c). Overall, this analysis points out that *freq*_*ext* and *freq*_*fix*_*avr* are relatively well explained while *gen*_*fix*_*avr* is more stochastic.

**Figure 3.**
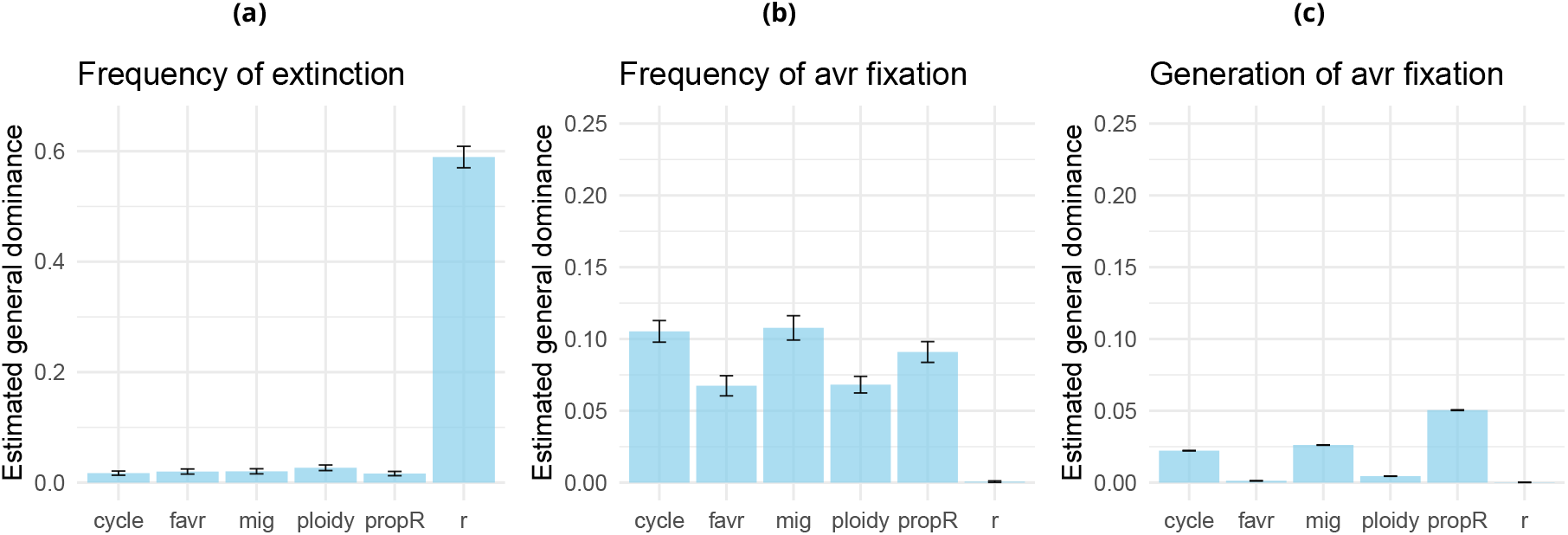
Estimated general dominance of each predictor calculated from general linear models applied to three output variables of the random simulation design: the frequency of extinction of population, the frequency of fixation of the *avr* allele among replicates with surviving populations, and the generation of fixation of the *avr* allele. For each predictor the general dominance was estimated from bootstrap average values with the corresponding standard error for 100 bootstrap resamples.

### Patterns of virulence fixation

Three different and exclusive equilibria are observed: the extinction of the pathogen population, the fixation of the *avr* allele and the fixation of the *Avr* allele. The frequencies of these outputs among replicates are represented depending on *f*_*avr*_, *r, propR* and *Cycle*, for haploids and diploids (Figure 4, Annex B Figure S5). For both ploidy levels, there is an increase in the frequency of fixation of the *avr* allele with the increase in *f*_*avr*_ and *r*. This representation also highlights the existence of a growth rate threshold *r*_0_ above which there is fixation of either the *avr* allele or the *Avr* allele, and below which there is instead either fixation of the *avr* allele or extinction of the population. In other words, for a growth rate below *r*_0_ the pathogen population only survives when virulent individuals invade the R compartment, which corresponds to evolutionary rescue. Evolutionary rescue is particularly noticeable for the life cycle with host alternation because in this case, *r*_0_ increases with *propR*.

**Figure 4.**
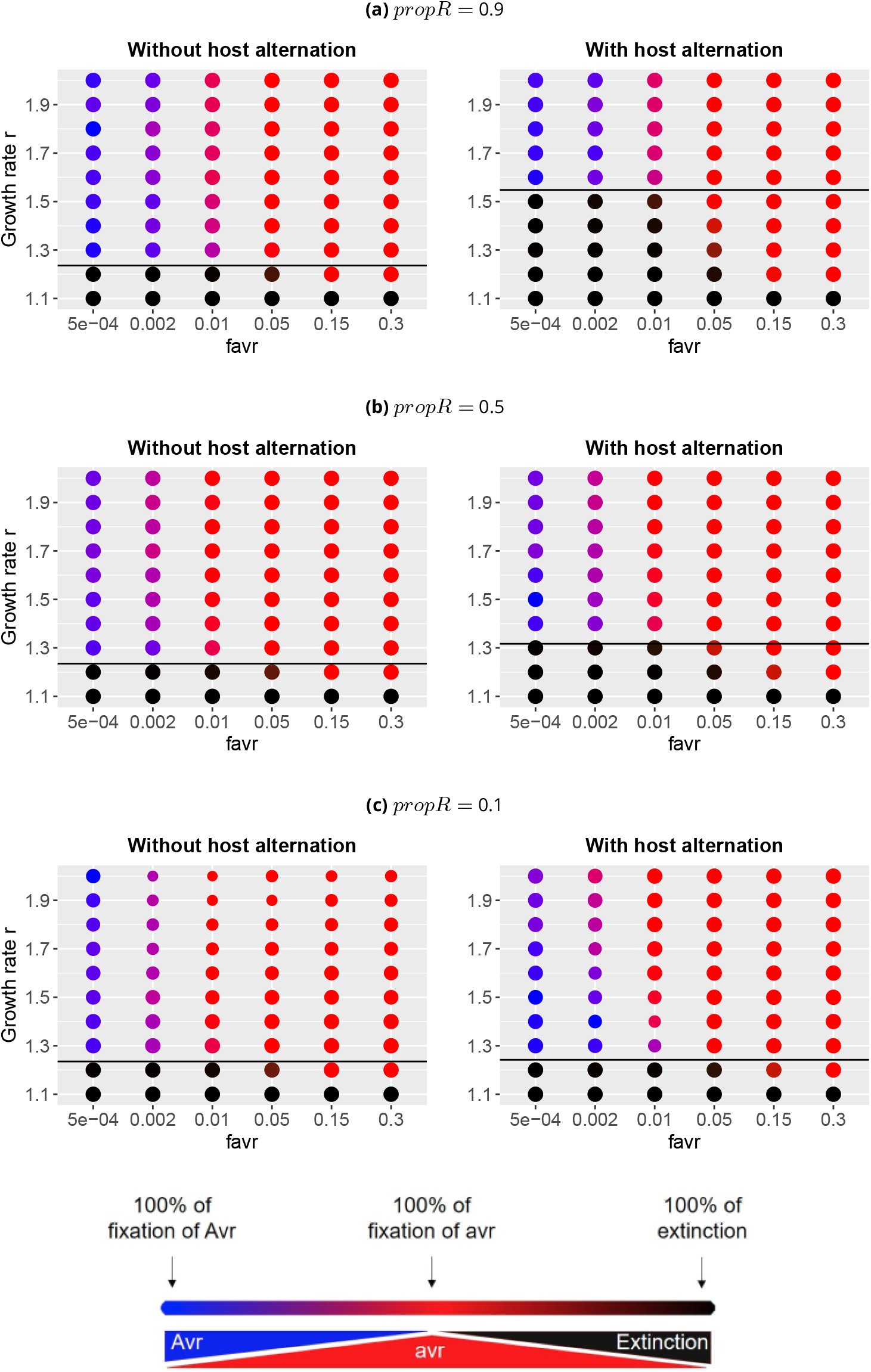
For diploid pathogens, representation of the frequencies of population extinction or fixation of alleles *Avr* or *avr* for each combination of four parameters: *f*_*avr*_, *r, propR* and *cycle*, with *mig* = 0.05. On each graph the black line corresponds to the calculated value of the growth rate threshold *r*_0_ below which the population dies if it does not expand to the R compartment. The surface of each plotted result is proportional to the number of simulations, among the 100 replicates, for which an equilibrium was reached at the end of the 1100 generations simulated.

Above *r*_0_, we observe a gradient between the predominant fixation of the two alleles depending of *f*_*avr*_, with slightly different patterns influenced by *propR, Cycle* and *r* for diploids (Figure 4). The influence of the life cycle on the fixation pattern is the most noticeable for low values of *propR* and *r*. The frequency of fixation of the *avr* allele appears maximal for intermediate values of *propR*.

To examine further the influence of *propR* on the probability of fixation of the *avr* allele, we plotted the evolution of the frequency of fixation of the *avr* allele among all replicates of the regular simulation design depending on *propR* and *r* for a fixed value of *f*_*avr*_ (Figure 5). For haploid individuals with host alternation, the frequency of fixation of the *avr* allele increases very slightly with *propR*. For haploids without host alternation, a plateau is observed for intermediate values of *propR*. For diploids, the distribution is shifted with a peak of maximal proportion of *avr* fixation for a slightly lower value of *propR*.

**Figure 5.**
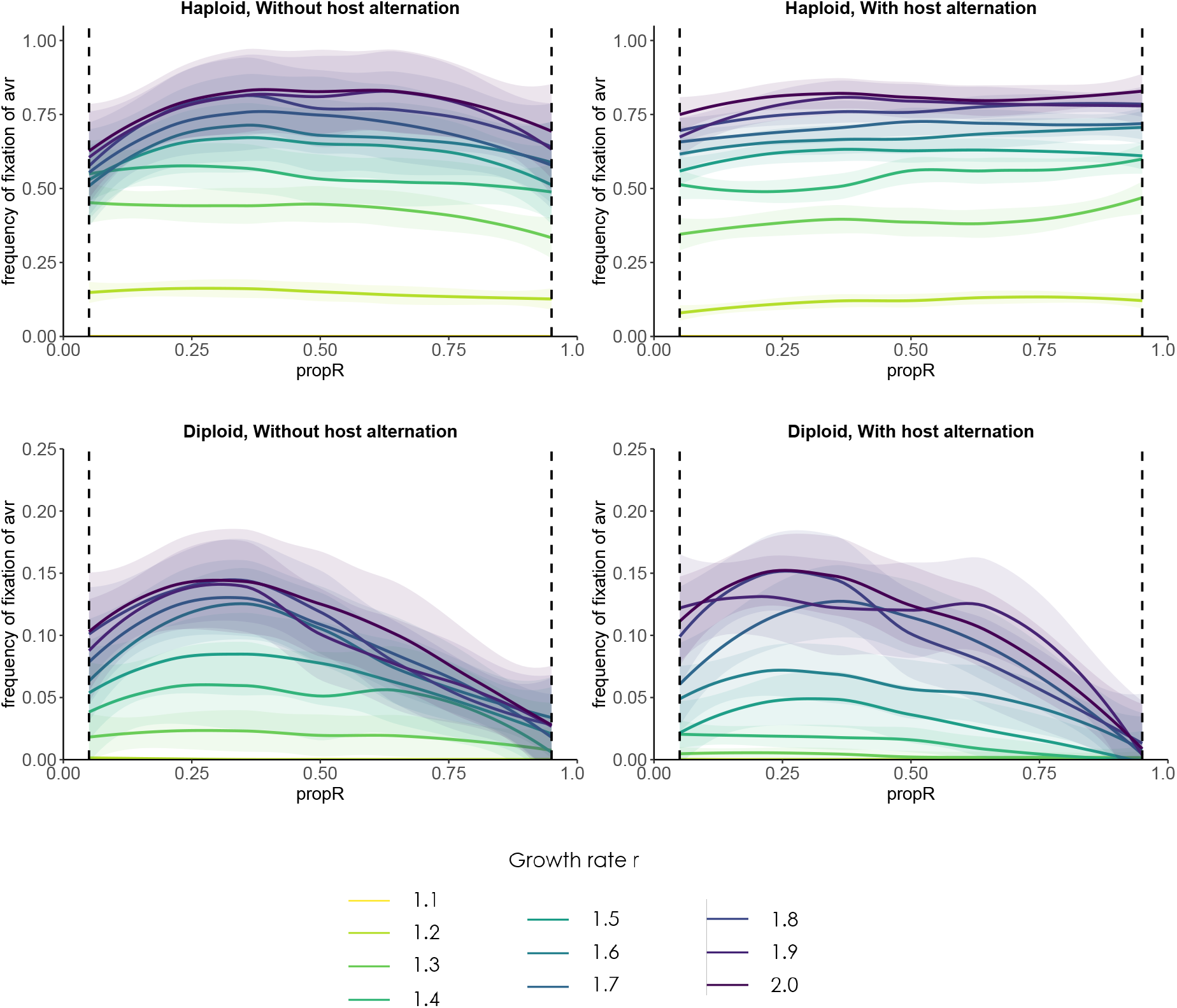
Evolution of the frequency of fixation of the *avr* allele depending on *propR* for *r* varying between 1.1 and 2.0, with *f*_*avr*_ = 0.0005. Simulations were performed without and with host alternation, for haploid and diploid individuals, with 100 replicates for each combination of parameters. The plotted results correspond to the local regression on the frequency of fixation of the *avr* allele with the 95% confidence intervals associated with each regression. The vertical dotted lines correspond to the bounds of simulated values of *propR* for this regular experimental design.

### Variations in the speed of virulence spread

To analyse in more details the dynamics of virulence spread, we examine two time points, reflecting two measures of resistance durability: the invasion of the R compartment and the resistance breakdown event. Invasion and resistance breakdown were defined as the first year when at least 1‰ and 1% of the resistant compartment were occupied by pathogens, respectively. Distributions of these two measures of resistance durability were plotted for three values of *propR*, with and without host alternation, only for the replicates for which we observed eventually a fixation of the *avr* allele. To broaden the picture, we monitored also the evolutionary trajectory of the *avr* allele from the invasion of the R compartment. The dynamics of invasion is mainly driven by the ploidy level and the dynamics of virulence spread is mainly driven by *propR* (Annex B Figure S3).

For haploid individuals resistance breakdown occur very rapidly: during the first or second year of simulation, regardless of the life cycle (figure not shown). There is a small delay in the time of the resistance breakdown with the increase in *propR*, especially without host alternation.

For diploid individuals, we observed longer periods before invasion and resistance breakdown and higher kurtosis. Assuming a strong migration rate, there are few differences between life cycles on the time of invasion. Both life cycles display an increase in kurtosis that goes hand in hand with the increase in *propR*. Without host alternation, distributions of the year of resistance breakdown and invasion time are similar, but with a one-year lag. Conversely, with host alternation, there are strong changes in the distributions that flatten out when considering the year of resistance breakdown (Figure 6). Assuming a low migration rate (i.e. for telluric pathogens), distributions of the year of invasion and resistance breakdown remain unchanged with host alternation, while these distributions flatten out considerably without host alternation (Annex B Figure S4). Note that in both migration regimes, with host alternation we observe a bimodal distribution of invasion year for *propR* = 0.9, with many invasion events in the first year of simulations. Early invasion events result from the initial redistribution of pathogen individuals following sexual reproduction on the alternate host: it is all the more likely that a virulent individual arrives on the R compartment the more predominant it is in the landscape.

**Figure 6.**
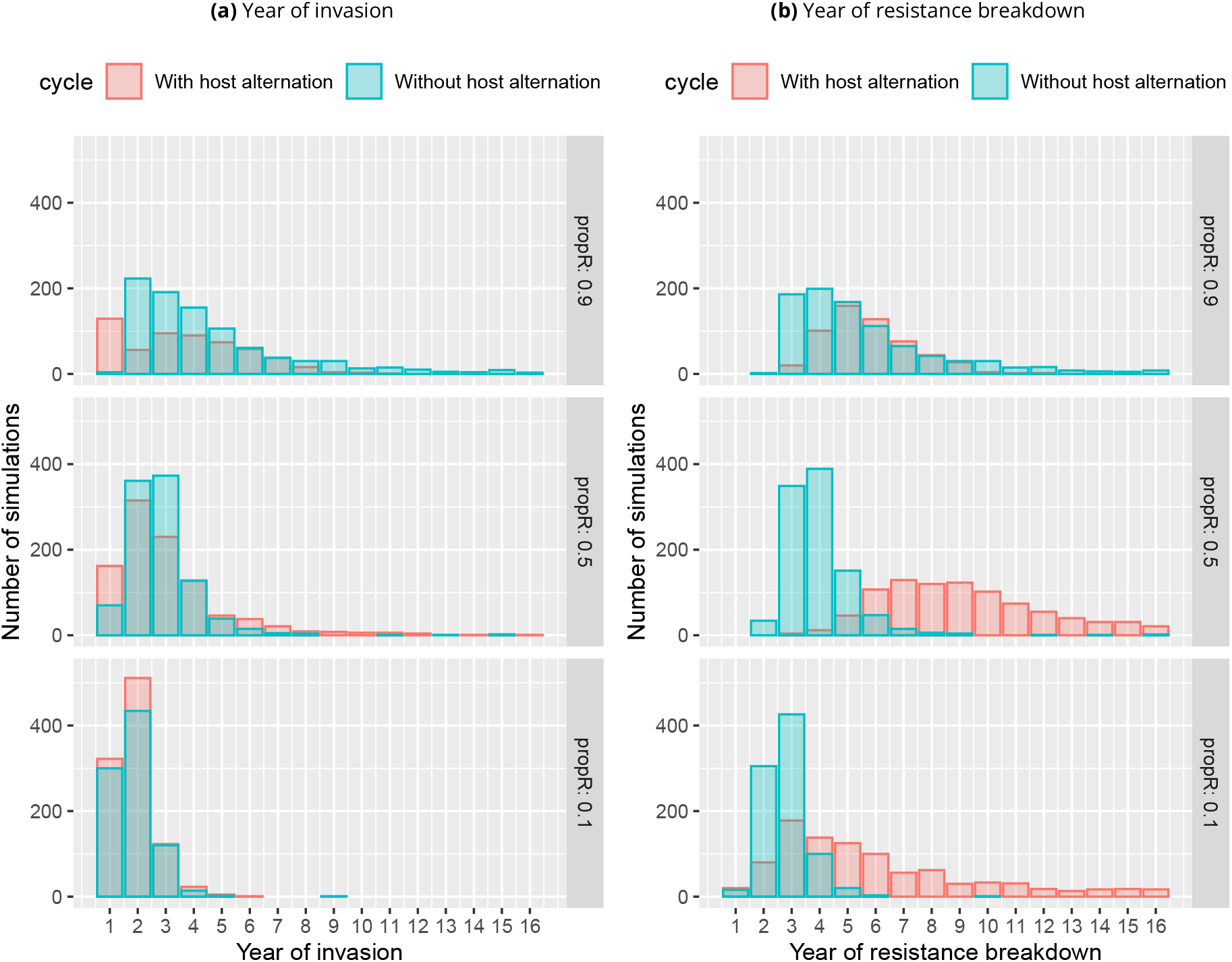
Histograms of (a) the year of invasion and (b) the year of resistance breakdown depending on the life cycle of the diploid pathogen, for three values of *propR*. Simulations were performed with *f*_*avr*_ = 0.02, *mig* = 0.05, and *r* = 1.5. The plotted results were obtained from the restricted simulation design, and correspond to all simulations among the 1 000 replicates per combinations for which we observed a fixation of the *avr* allele in the population.

In a last step, the evolution of the frequency of the *avr* allele in the population was studied along with the evolution of the population sizes through generations, from the invasion (Figure 7, see Annex B Figure S6 for haploids). For both life cycles, we observe an increase in the speed of fixation of the *avr* allele with the increase in *propR*. The median speed of fixation is higher with host alternation for haploids, and highly dependant of *propR* without host alternation for diploids (Annex B Figure S3). We also observe differences in stochasticity levels depending on the ploidy and the life cycle. For haploids, the dynamics of virulence fixation is almost deterministic. For diploids, the dynamics are more variable, with a highly stochastic behaviour for the life cycle with host alternation. Moreover, the results of GLM, the dominance analysis and the comparison of both figures show that independently of *propR* and of the life cycle, the increase in the proportion of *avr* allele is faster for haploid than for diploid individuals.

**Figure 7.**
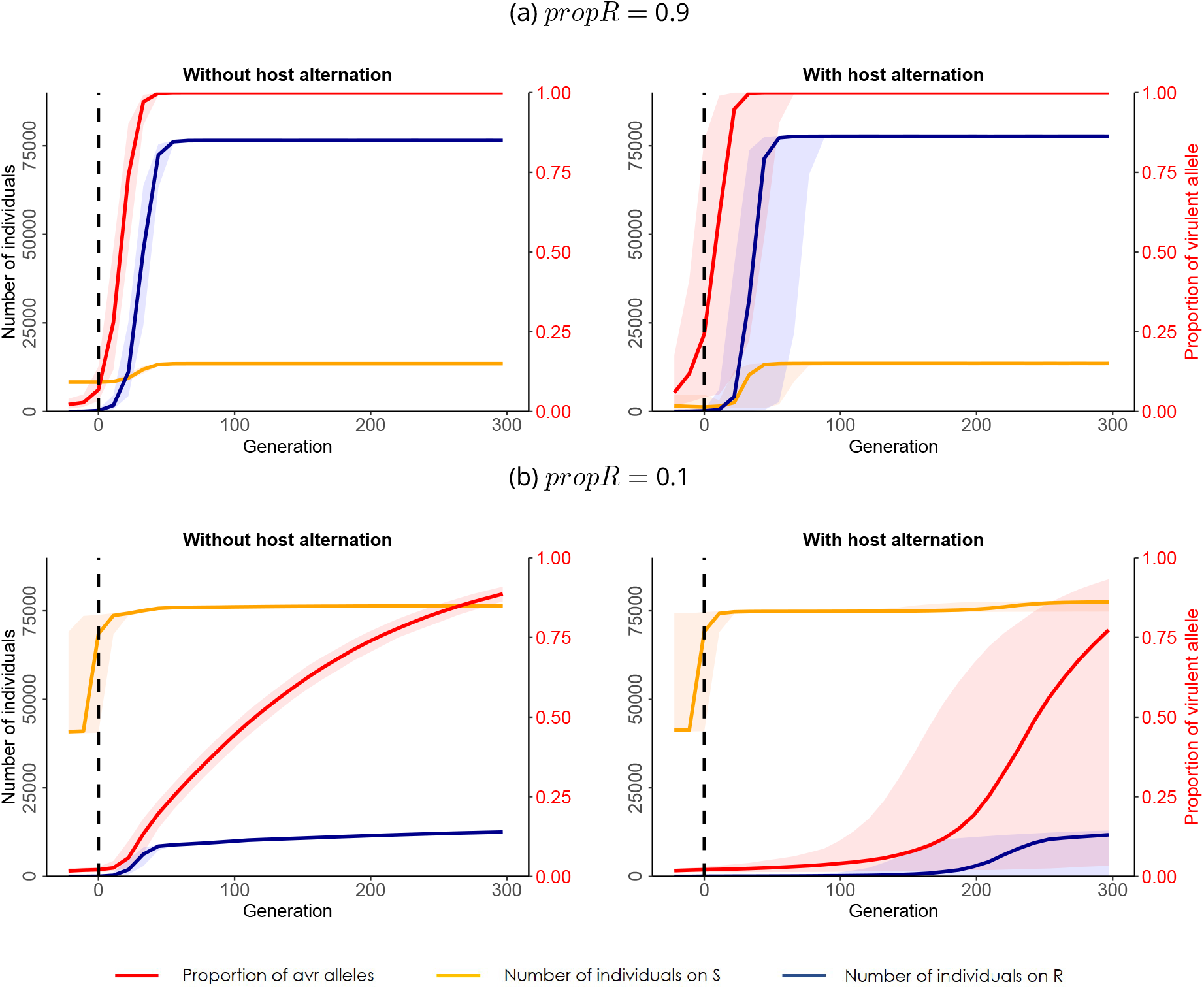
For diploid pathogens, evolution of the population size in the R and S compartments (on the left scale) and of the frequency or *avr* alleles in the S compartment (on the right scale) through generations. Simulations were performed without and with host alternation, for two values of *propR*: (a) 0.1 and (b) 0.9, with *f*_*avr*_ = 0.02, *mig* = 0.05, and *r* = 1.5. For each simulation, generation 0 corresponds to the generation at which the invasion occurred. For each combination of parameters, simulations were performed on 1 000 replicates. The plotted results correspond to the median results (frequency of *avr* alleles or population size) for all simulations among the 1 000 replicates for which we observed a fixation of the *avr* allele in the population. Coloured intervals correspond to the 95% confidence intervals.

Focusing on diploid simulations, Figure 7.b highlights the existence of an evolutionary rescue effect, for the life cycle with host alternation and a high value of *propR*. The median population size decreases through time and almost reached 0 before the 20th generation following the invasion, when an increase in the proportion of *avr* alleles lead to an increase in the population size in the R compartment, followed by an increase in the population size of the S compartment, preventing the extinction of the population.

Interestingly, the speed of invasion is mainly driven by the ploidy, while the speed of fixation of the *avr* allele from the invasion is mainly driven by the landscape composition *propR* (Annex B Figure S3).

## Discussion

### Deep impact of the ploidy on resistance durability

Lof et al. (2017) demonstrated that the pre-existence of virulent alleles in the pathogen population could greatly diminish resistance durability. In the present study, we varied the initial frequency of *avr* alleles in the population but focused only on cases where this allele was already present at the beginning of the simulation, which corresponds to standing genetic variation (Alexander et al., 2014; Barrett et al., 2008). Our results illustrated a strong positive relationship between the initial frequency of virulent alleles and the probability of invasion and resistance breakdown. For haploid individuals, we found no stochasticity in the time of occurrence of the invasion, which occurred in the first year of the simulation for all replicates. Thus, for simulations with haploid pathogens, almost as soon as one virulent individual invaded the resistant compartment, it was selected and the resistance breakdown occurred. This result explains why a lot of models on haploid individuals focus on the probability of apparition of the first virulent individual, in particular by mutation (Fabre et al., 2015; Papaïx et al., 2015). Our results for simulations with haploid pathogens also highlighted low stochasticity in the increase in the proportion of virulent alleles in the population after the invasion. The results obtained with haploids were consistent with previous studies on resistance durability (Pacilly et al., 2018), which permitted us to consider this model as the reference model, in order to study the influence of the diploid state on the system dynamics. Diploid individuals, however, display strongly different evolutionary trajectories. In particular, we observed higher stochasticities in the evolution of the virulent allele frequency, both before and after the resistance breakdown. This is mainly caused by the recessivity of the *avr* allele, according to the gene-for-gene model. Before the invasion, the *avr* allele is rare and mostly at the heterozygous state, hence leading to phenotypically avirulent individuals. Therefore, the *avr* allele is poorly selected, and variations in its frequency are mostly driven by genetic drift, which induces high stochasticity among replicates. This effect should be strengthened in species with small effective population sizes, such as in cyst nematodes (Gracianne et al., 2016; Montarry et al., 2019).

As a consequence, we observed lower frequencies of *avr* fixation and higher extinction rates for diploid individuals, independently of the life cycle and the host deployment strategy. Moreover, simulations with diploid pathogens resulted in lower speeds of fixation of the virulent allele, that is, higher resistance durability. Because of the heterozygous *Avr*/*avr* state, *avr* alleles are not necessarily selected and their presence in the population does not inevitably lead to an immediate resistance breakdown, as observed for haploid individuals. Thus, besides its impact on the stochasticity of the results, the vulnerability of the virulent allele at the heterozygous state is also responsible for an increase in resistance durability. The impact of the landscape composition on resistance durability also differs with the ploidy. Consistently with the work of Van den Bosch and Gilligan (2003) and Pietravalle et al. (2006), for haploid individuals the increase in *propR* leads to a strong increase in the speed of fixation of the *avr* allele, thus decreasing the resistance durability. In all cases except for haploids with host alternation, this result was accompanied in the present study by a maximum frequency of *avr* fixation for intermediate values of *propR*. This non-linear relationship is similar to the one highlighted for haploids by Pacilly et al. (2018), and is caused by two distinct mechanisms. At low proportions of resistant hosts in the landscape, the selective pressure on the *avr* allele is sufficiently low to limit the increase in the virulence in the pathogen population. At high values of *propR*, the opposition between selection and drift is magnified: on one hand the selective pressure is high and imposes a rapid pace of adaptation; on the other hand the small size of the S compartment increases genetic drift and the risk of extinction of the *avr* allele, provided that the R compartment is not invaded. Hence, in most cases the virulent allele is lost if it does not spread quickly enough in the population: either *r > r*_0_ which lead to a fixation of the avirulent *Avr* allele, either *r < r*_0_ and the population goes extinct. Therefore, it would be possible to reduce the probability of invasion for diploid pathogens with either very low or very high proportions of resistant hosts in the landscape. However the increase in the proportion of resistant hosts is at the risk of weaker resistance durability: if the resistance breakdown occurs, it occurs more rapidly. For haploid pathogens with host alternation, we observed an almost constant and slightly increasing frequency of fixation of *avr* with the increase in *propR*. In this case and contrary to diploids with host alternation, the increase in the proportion of *avr* alleles on the resistant host is not counteracted by the allele reshuffling during the sexual reproduction event on the alternate host. For diploids, this allelic reshuffling causes a rise in the number of phenotypically avirulent *Avr*/*avr* heterozygous individuals, which will die if the redistribution following the sexual reproduction lead them on the resistant host. This can result in a drastic drop in the *avr* allele proportion while for haploids, the proportion of resistant hosts in the landscape does not increase the mortality rate of individuals carrying the virulent allele, because these haploid individuals are necessarily surviving on the resistant host.

### Life cycles impose different selection regimes and lead to contrasted evolutionary trajectories

The two different life cycles considered in this study - with or without host alternation - can be assimilated to hard and soft selection respectively (Bugge Christiansen, 1975; Wallace, 1975). Soft selection is expected to protect polymorphims, and hence promote local adaptation, while hard selection resembles an all or nothing game, that is to adapt to the encountered environment or to perish. Here host alternation can lead to faster evolution of allelic frequencies, with higher speeds of virulence fixation, especially for high values of *propR*. Without host alternation on the contrary, the evolution of virulence alleles are buffered, which result in more constrained dynamics. The increase in gene flow resulting from host alternation limits natural selection and local adaptation (Lenormand, 2002), especially because of the dispersal of non-adapted individuals on resistant hosts. The life cycle with host alternation is thus characterized by higher probabilities of population extinctions, and strong dependency of the growth rate threshold *r*_0_ and the landscape composition *propR*. For diploids with host alternation, contrary to the local adaptation on each compartment without host alternation, the forced allelic reshuffling on the alternate host is responsible for the increase in the number of *Avr*/*avr* heterozygous individuals. Because the *avr* allele is recessive, a large proportion of these newly-produced individuals die from the redistribution on the resistant compartment following the sexual reproduction. Noticeably, the reduction in virulence fixation at high proportions of resistant host discussed above hence results from two mechanisms in diploids: impediment of local adaptation without host alternation or increase in selective pressure with host alternation.

For diploid individuals, we also observed contrasting patterns of variation in the evolution of the *avr* allele frequency before and after the invasion, depending on the life cycle. The time of invasion is more stochastic without host alternation, while the speed of increase in the *avr* allele frequency from the invasion is far more stochastic with host alternation. The first step relies essentially on the probability of encounter between a virulent individual and a resistant host. Without host alternation the encounter is restrained to the probability that a virulent individual migrates from the susceptible to the resistant host during the vegetative season. The host alternation reinforces gene-flow, with the annual migration event that redistributes pathogen individuals among all host plants, thereby favoring the encounter. Interestingly, in the case of host alternation, early infections of resistant host (invasion) does not readily translate into population establishment on that host (disease outbreak leading to resistance breakdown). At the end of the vegetative season and initial invasion, the sexual reproduction on alternate host reshuffles allele frequencies, and thus breaks virulent (homozygous) individuals into mostly avirulent (heterozygous) individuals. These up and down selection phases amplify the effect of genetic drift and lead to nearly chaotic evolutionary trajectories, resulting in a resistance durability all the more difficult to predict. Without host alternation, virulent individuals mate with each others and the homozygous state is sheltered, which results in a strict time lag between initial invasion and population outbreak. Overall our model is in accordance with the framework proposed by McDonald and Linde (2002) which highlights the importance of gene flow as an impediment to resistance durability. Our analysis completes this framework, taking into account the variation in life cycles.

The life cycle also plays a role in the frequency of observation of evolutionary rescue effects. Carolan et al. (2017) highlighted the impact of the growth rate of the pathogen on resistance durability, by presenting the limitation of the growth rate as a mean to increase resistance durability. In accordance with this study, we displayed a growth rate threshold *r*_0_ below which the pathogen population goes extinct if it does not invade the resistant compartment. Hence, for a growth rate below *r*_0_, the genetic adaptation of the pathogen population is the only way for the population to survive, which is a classical example of evolutionary rescue. In the current study, *r*_0_, and thus the observation range of evolutionary rescue, is highly dependent on the life cycle. Without host alternation, the redistribution of individuals between compartments and the mortality rate is limited, which leads to a quite low *r*_0_, independently of the proportion of resistant hosts in the landscape. With host alternation, however, the redistribution event occurring each year from the alternate host to the S and R compartments leads to a strong dependence of *r*_0_ on the landscape composition, with an increase in the observation range of evolutionary rescue with the proportion of resistant hosts.

## Conclusion

Short-term epidemiological control is predicted to be optimal for a landscape composed of a high proportion of resistant hosts in a low degree of spatial aggregation (Holt and Chancellor, 1999; Papaïx et al., 2014, 2018; Skelsey et al., 2010). However other authors also highlighted that optimal resistance durability could be obtained by reducing the proportion of resistant hosts (Fabre et al., 2012; Papaïx et al., 2018; Pietravalle et al., 2006; Pink and Puddephat, 1999), thus minimising the selection pressure on the pathogen population. In the current study, the minimisation of the probability of fixation of the virulent allele for a diploid pathogen population was obtained either at very low or very high proportions of resistant hosts in the landscape. In cases where the population does not go extinct and the virulent allele increases in proportion, however, the proportion of resistant hosts in the landscape strongly impacts the speed of increase, and thus the resistance durability. Consistently with Van den Bosch and Gilligan (2003), we displayed that for a diploid pathogen population with standing genetic variation, the increase in the proportion of resistant hosts decreases resistance durability. In particular, with host alternation both the invasion and the fixation of the virulent allele in the population can occur very quickly. However, in such a case, the evolutionary trajectory of the virulent allele is all the more stochastic and durability is thus difficult to predict. Without host alternation (i.e. for the majority of pathogen species) early detection and population control measures would increase resistance durability. However such prophylactic measures are made all the more difficult by the strong unpredictability of the invasion date. For the few species with host alternation, a massive intervention could durably control a population of pathogens, such as the eradication of the alternate host species *Berberis vulgaris* for wheat stem rust control (Peterson, 2018). Overall, the high stochasticity of evolutionary trajectory impedes accurate forecasts of resistance durability for diploid organisms.

## Perspectives

In the current study, we considered a single qualitative resistance gene. The combination of several resistance genes is often studied and deployed to increase resistance durability (Djian-Caporalino et al., 2014; Djidjou-Demasse et al., 2017; Mundt, 2014; Rimbaud et al., 2018). These combinations of resistances can occur at the plant scale with multiple resistance genes (pyramiding) inside one host genotype, or at the landscape scale with multiple resistance deployments in time or space (Mundt, 2002; Van den Bosch and Gilligan, 2003). Some resistant cultivars progressively introduced in the landscape are composed of different combinations of qualitative resistance genes, resulting in an evolving selective pressure through time (Goyeau and Lannou, 2011). Building on our results, we can extrapolate on the impact of the ploidy and the life cycle on resistance durability for these different strategies of deployment. Hence, rotating cultures with different resistances would amount to force gene flow, favoring the encounter of pathogen individuals with new hosts without an actual migration. This would be comparable to the hard selection regime observed with host alternation, and we expect similar results. With the pyramiding of several resistance genes in same host, meanwhile, we would expect a higher short term efficiency than with a single resistance, but with a higher risk of rapid resistance breakdown by multi-virulent individuals due to the inducing of a strong selective pressure. This may especially be true for pathogens with host alternation because of the increased probability of mating between pathogens with different virulent profiles when they meet on the alternate host.

## Data accessibility

Python simulation code and results, as well as R script for result analyses are available on Zenodo repository: (DOI: 10.5281/zenodo.4892587).

## Acknowledgements

We warmly thank Josselin Montarry, Lydia Bousset, Jean-Paul Soularue, Frédéric Grognard, Bénédicte Fabre, Clémentine Louet, Pascal Frey, and Cyril Dutech for constructive comments on previous versions of the manuscript. We thank Frédéric Fabre for insightful discussions on sensibility analyses. We also thank Hirohisa Kishino, Loup Rimbaud and one anonymous reviewer for detailed comments and suggestions which helped improve the manuscript. Version 4 of this preprint has been peer-reviewed and recommended by Peer Community In Evolutionary Biology (https://doi.org/10.24072/pci.evolbiol.100131).

## Annex A: Sobol indices

Sensitivity analyses were performed with the calculation of Sobol indices (first-order, second-order and total-order) with the R package “sensitivity” (Iooss et al., 2021). Sobol indices were calculated to study the importance of each of the six parameters of interest on the output variable *freq*_*ext* only (figure S1). These calculations were based on the results issued from the random simulation design.

For the four remaining outputs (*freq*_*fix, gen*_*fix*, the year of occurrence of the invasion, and the time elapsed between the invasion and the fixation of the *avr* allele), only the simulations not leading to extinction were retained for the sensitivity analyses. Thus, the input combinations of parameters retained depended strongly on the results of the output variable *freq*_*ext*, the independence hypothesis of the input parameters were then not verified and Sobol indices could not be calculated for these four remaining output variables. Further analyses would be necessary to disentangle the effect of each parameter of interest on these remaining output variables.

**Figure S1.**
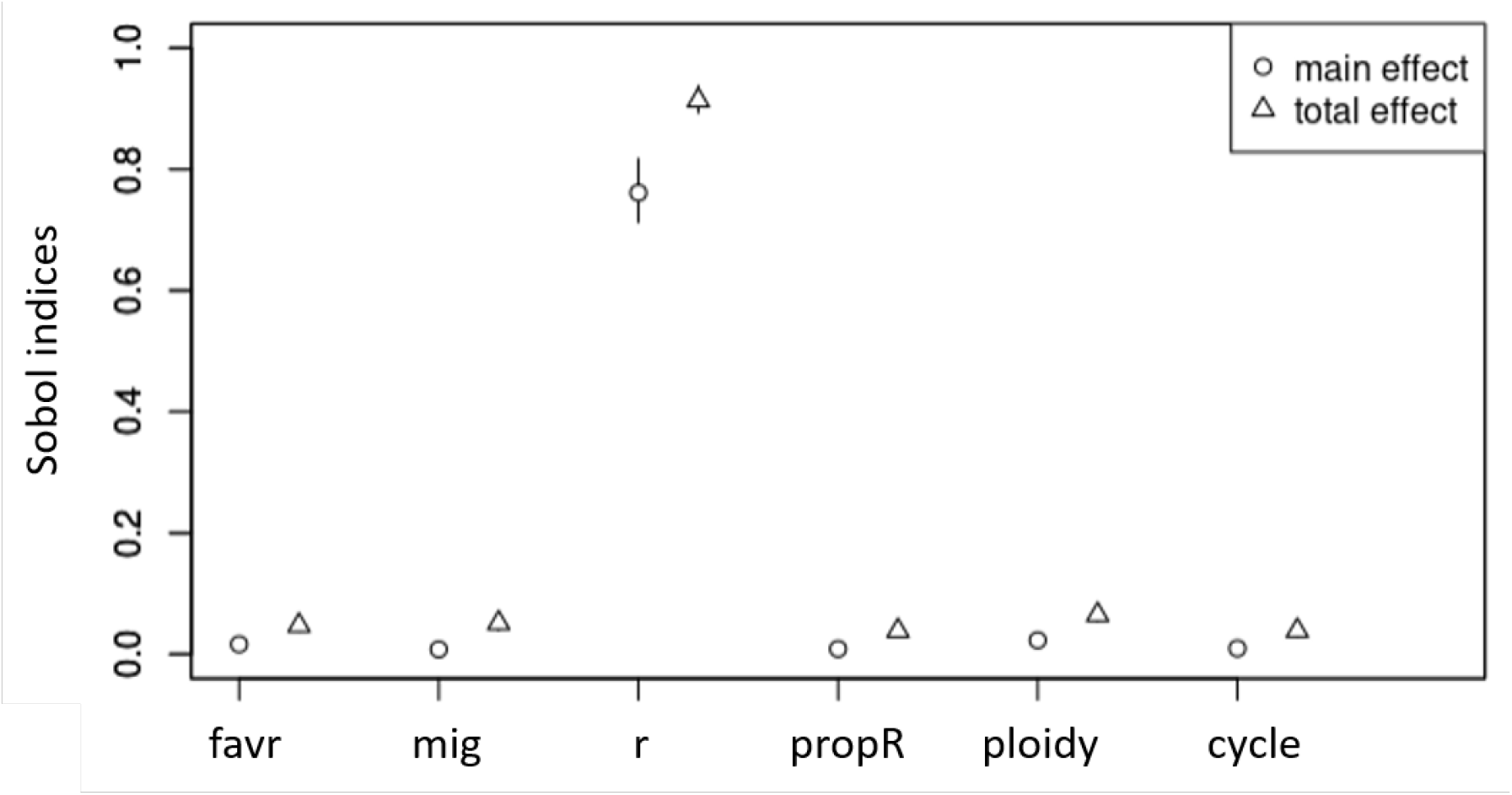
Sobol indices to evaluate the influence of six variables of interest on the frequency of extinctions among simulations. Main effect correspond to first-order Sobol indices, and total effect correspond to total-order Sobol indices.

## Annex B: Supplementary figures

**Figure S2.**
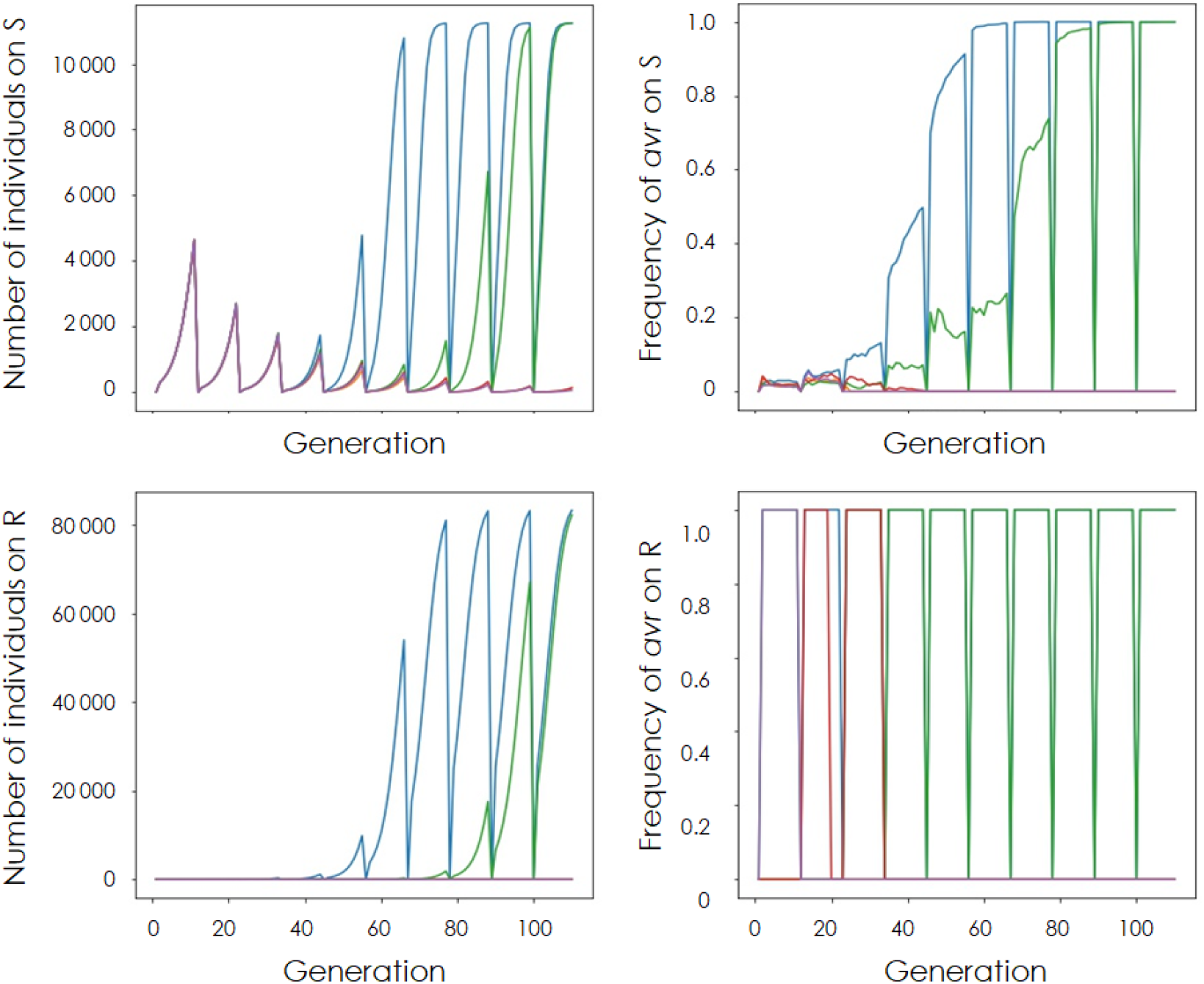
Example of the simulated populations demographic (on the left) and virulent allele frequency (on the right) dynamics through time on S and R compartments. The model was run for four replicates with diploid individuals and host alternation, *propR* = 0.9, *r* = 1.5, *f*_*avr*_ = 0.025, and *mig* = 0.05. Each color represents a distinct replicate.

**Figure S3.**
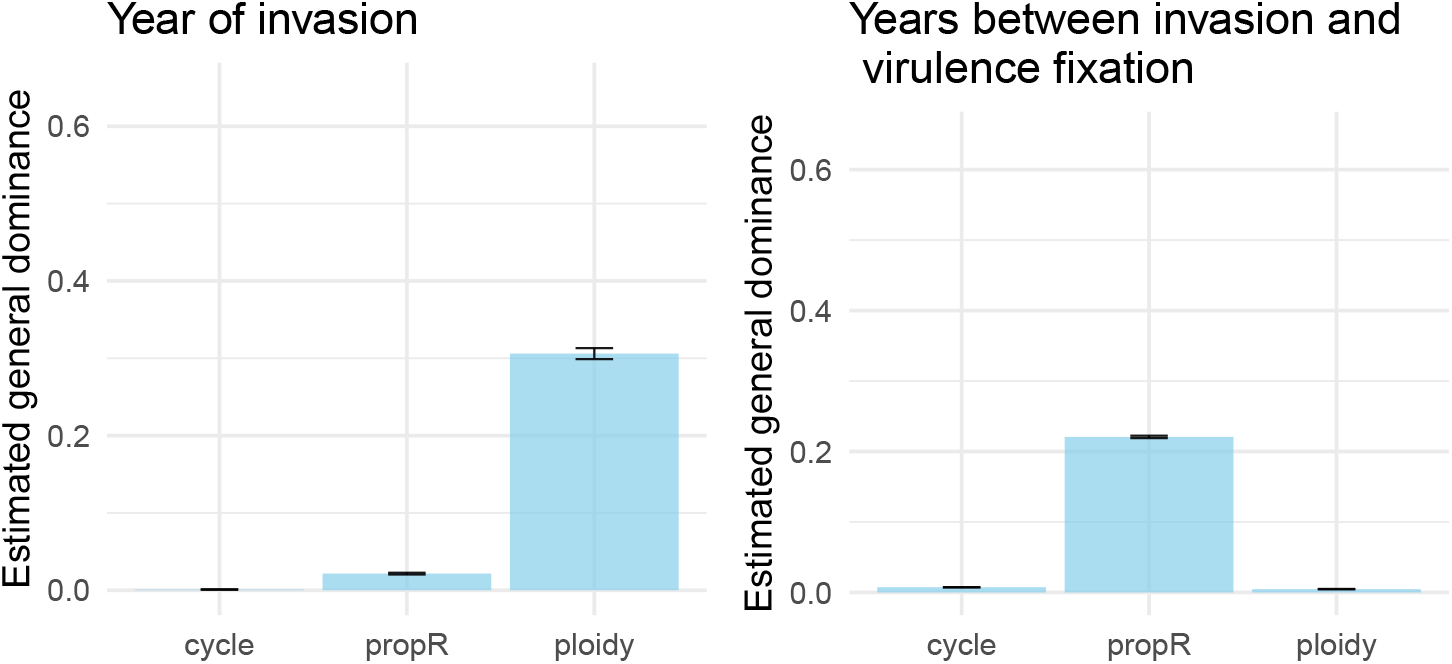
Estimated general dominance of each predictor calculated from general linear models applied to two output variables of the restricted simulation design: the year of invasion and the time elapsed between the invasion and the fixation of the *avr* allele. For each predictor the general dominance was estimated from bootstrap average values with the corresponding standard error for 100 bootstrap resamples.

**Figure S4.**
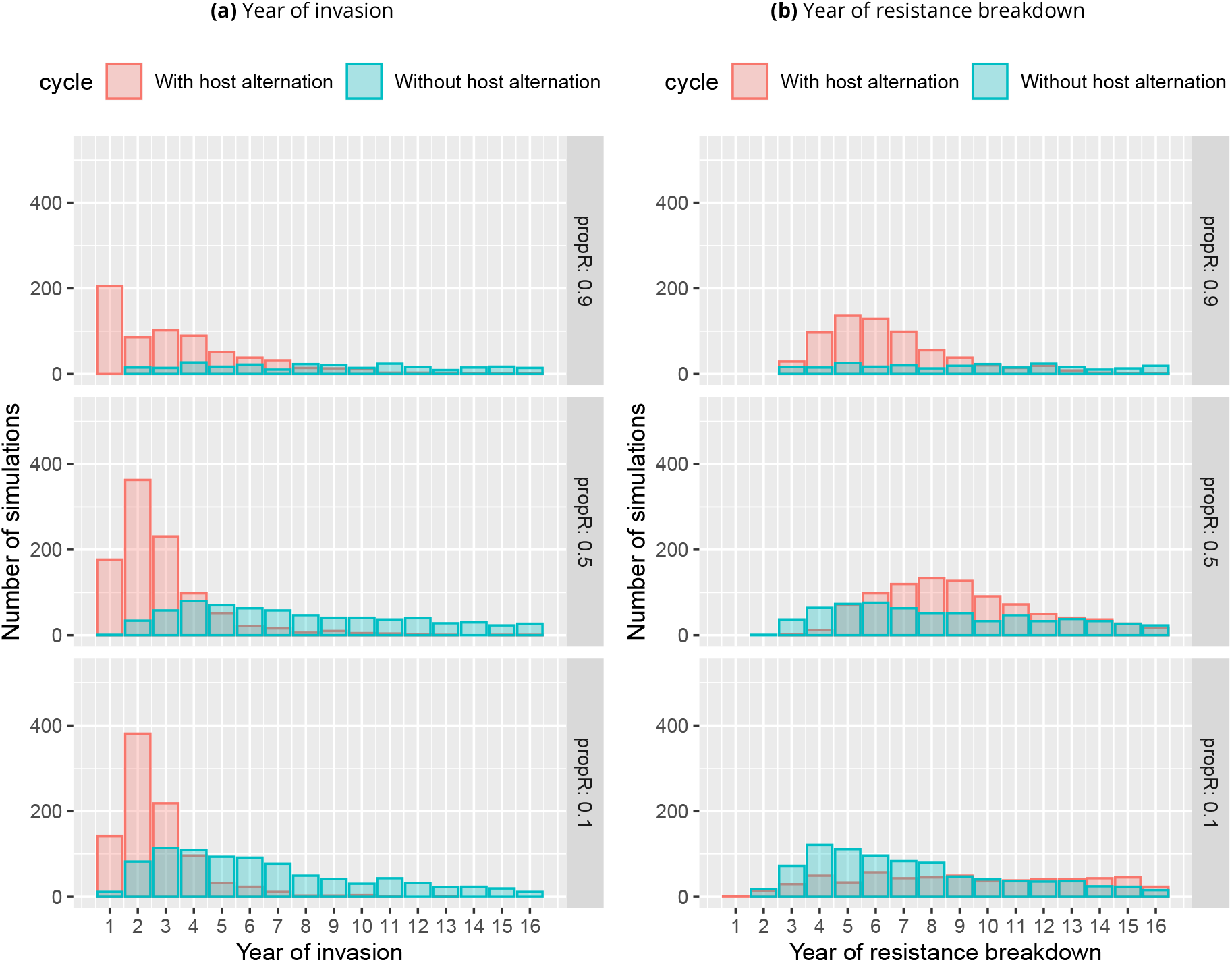
Histograms of (a) the year of invasion and (b) the year of resistance breakdown depending on the life cycle of the diploid pathogen, for three values of *propR*. Simulations were performed with *f*_*avr*_ = 0.02, *mig* = 0.001, and *r* = 1.5. The plotted results were obtained from the restricted simulation design, and correspond to all simulations among the 1 000 replicates per combinations for which at least 80% of the R compartment is occupied at the end of the simulation.

**Figure S5.**
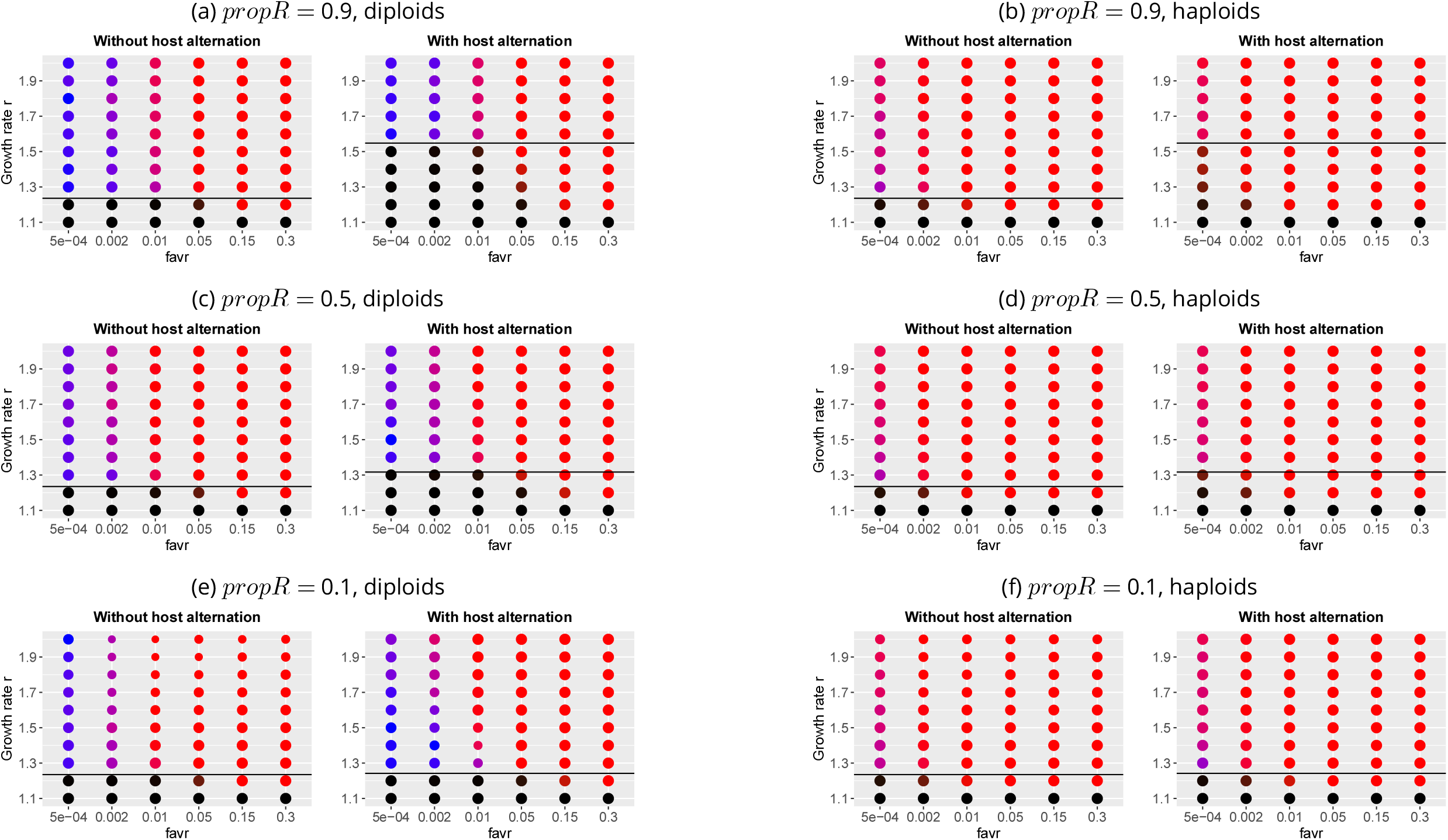
Representation of the frequencies of population extinction or fixation of alleles *Avr* or *avr* for each combination of five parameters: *f*_*avr*_, *r, propR, ploidy* and *cycle*, with *mig* = 0.05. On each graph the black line corresponds to the calculated value of the growth rate threshold *r*_0_ below which the population dies if it does not expand to the R compartment. The surface of each plotted result is proportional to the number of simulations, among the 100 replicates, for which an equilibrium was reached at the end of the 1100 generations simulated. In colored dots, red corresponds to the fixation of the *avr* allele, blue to the fixation of the *Avr* allele, and black to the extinction of the population. Dot color shades indicate simulation results among replicates. The left part ((a), (c), and (e)) corresponds to Figure 4, presented in the main document.

**Figure S6.**
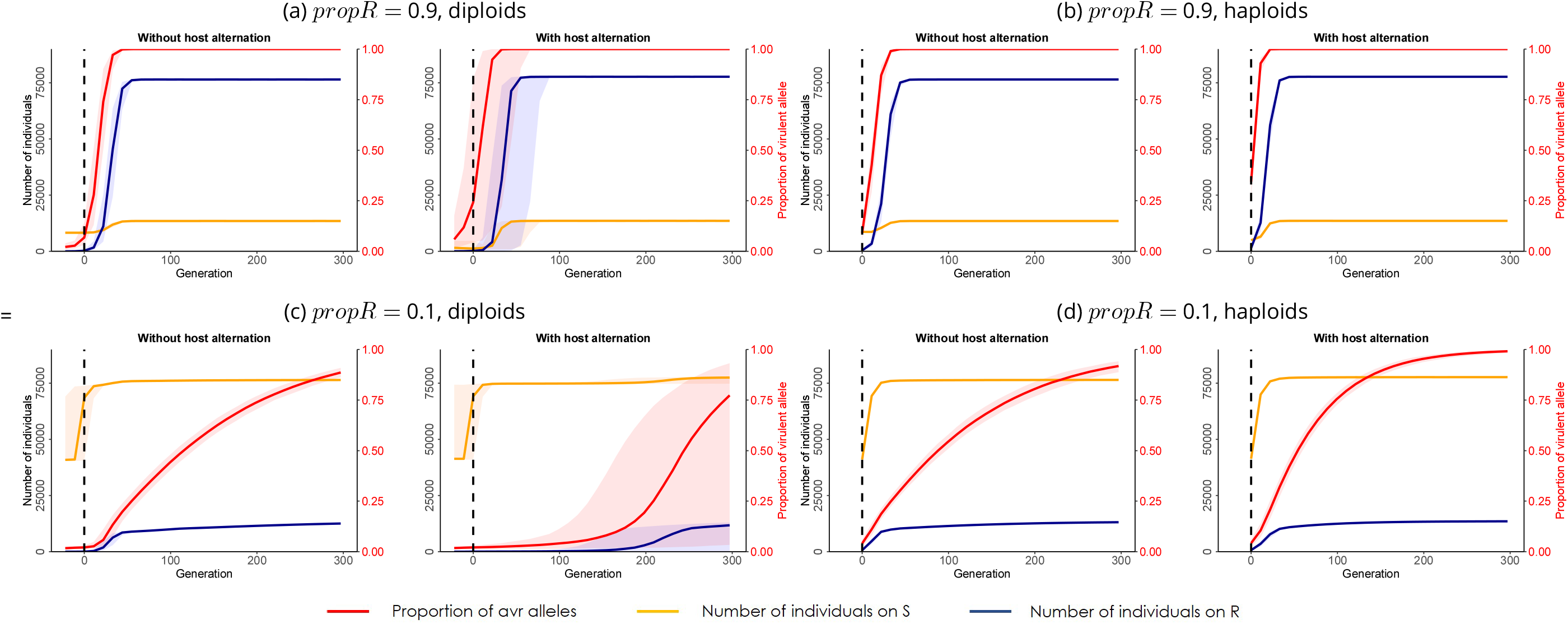
Evolution of the population size in the R and S compartments (on the left scale) and of the frequency or *avr* alleles in the S compartment (on the right scale) through generations. Simulations were performed without and with host alternation, for (a, c) diploid and (b, d) haploid pathogens, for two values of *propR*: (a, b) 0.9 and (c, d) 0.1, with *f*_*avr*_ = 0.02, *mig* = 0.05, and *r* = 1.5. For each simulation, generation 0 corresponds to the generation at which the invasion occurred. For each combination of parameters, simulations were performed on 1 000 replicates. The plotted results correspond to the median results (frequency of *avr* alleles or population size) for all simulations among the 1 000 replicates for which we observed a fixation of the *avr* allele in the population. Coloured intervals correspond to the 95% confidence intervals. The left part ((a) and (c)) corresponds to Figure 7, presented in the main document.

